# aiAtlas: High-Fidelity Cell Simulations of Genetic Perturbations in Rare Diseases and Cancers

**DOI:** 10.1101/2025.10.20.683396

**Authors:** Wayne R Danter

## Abstract

**Background:** Linking genetic perturbations to cellular phenotypes remains a central challenge in translational biology. Experimental iPSC and organoid models are powerful but constrained by scalability, variability, and difficulty modeling rare or polygenic states.

**Methods:** We developed **aiAtlas v1.2**, a high-fidelity simulation platform that integrates Large Concept Model (LCM) logic with aiPSC-derived modeling. We evaluated **136 virtual cell lines** spanning wild-type, single-mutation, multiple-mutation, human tumor-derived, and gene-fusion cohorts. **Twenty-five** features covering DNA damage/repair, replication stress, epigenetic remodeling, pluripotency, and stress responses were quantified. Statistical analysis used the **Mann–Whitney U test** with **Bonferroni correction, Hodges–Lehmann estimators (HLE)** for median differences, and **Cliff’s delta** effect sizes with **bootstrap 95% confidence intervals**. Robustness measures included early stopping, bagging, and 5-fold cross-validation.

**Results:** aiAtlas v1.2 **reliably separated wild-type and mutant cohorts**, revealing consistent disruptions in DNA damage accumulation, replication stress, epigenetic dysfunction, and loss of pluripotency, while identifying **stable features** (e.g., core nucleotide-excision repair processes and selected apoptosis measures). Subgroup analyses showed shared systemic effects and **context-specific vulnerabilities**: single mutations frequently produced measurable divergence; multiple mutations amplified instability; tumor-derived and gene-fusion lines yielded distinct but partially overlapping phenotypes. **Large effect sizes** (Cliff’s δ) with **narrow bootstrap CIs** supported reproducibility across cohorts.

**Conclusions/Impact:** aiAtlas v1.2 provides a **robust virtual subject framework** that uses aiCRISPR-Like (aiCRISPRL) virtual gene editing system that complements wet-lab CRISPR models by scaling to diverse genomic contexts and highlighting both disruption and stability. The platform can guide therapeutic prioritization, gene-editing strategy design, and regulatory innovation consistent with the **FDA Modernization Act 2.0**, accelerating therapy development in rare diseases and cancer.

**Significance Statement:** aiAtlas introduces a scalable, reliable simulation framework that integrates advanced large concept model (LCM) logic with iPSC-derived cellular modeling. aiAtlas overcomes major limitations of experimental systems by capturing both broad and subgroup-specific phenotypic divergence across single mutations, multiple mutations, tumor-derived cell lines, and gene fusions. This reliable platform establishes a new opportunity for rare diseases and cancer modeling, offering reproducible insights that can accelerate discovery and translational applications where traditional wet-lab approaches are impractical.

Furthermore, in situations where the target mutational profile has been defined but no cellular models yet exist, aiAtlas can quickly generate custom virtual cell lines that accurately reproduce the corresponding genomic and phenotypic features.

## Introduction

Understanding how genetic alterations drive disease phenotypes at the cellular level remains one of the fundamental challenges in biomedical research. While induced pluripotent stem cells (iPSCs) and established human cell lines have provided powerful experimental systems, they remain constrained by cost, scalability, and an inability to fully capture the systemic consequences of complex genomic states [1,2]. These limitations are particularly important in the context of rare and ultra-rare diseases, where patient material is often scarce [3], and in cancers, where multiple mutations and gene fusions often interact in unpredictable ways [4,5].

While evolving computational models attempt to bridge this gap, most current approaches still lack the fidelity to reproduce multi-pathway interactions at cellular and systems levels [6]. There remains an unmet need for a platform that can integrate genomic, epigenomic, and cellular features to produce more physiologically realistic, and testable predictions.

We recently developed aiAtlas, a simulation-based platform built on large concept model (LCM) logic and aiPSC modeling, designed to simulate cellular states and predict phenotypic divergence arising from a broad array of genetic perturbations [6,7]. Large Concept Models (LCMs) represent a significant advancement over traditional Fuzzy Cognitive Maps (FCMs). While FCMs are constrained to explicit networks of weighted causal relationships defined between user-specified concepts (nodes), LCMs operate in a high-dimensional concept space that allows for abstraction, generalization, and the emergence of new properties beyond predefined variables. LCMs represent a powerful AI architecture that extends beyond basic causal linkages to capture the broader conceptual dynamics of complex systems, making them a powerful evolution of FCM logic for high-fidelity biological simulation.

In Part 1, we evaluate the ability of aiAtlas to reliably distinguish wild type (WT) aiPSCs from a diverse population of mutant cells, capturing major shifts in cell cycle regulation, DNA repair, epigenetic remodeling, pluripotency, and the hallmarks of cancers.

In Part 2 of this study, we extend this framework to clinically relevant subgroups, each representing a distinct biological context including single gene mutations, multiple mutations, human tumor-derived cell lines, and gene fusions. By systematically comparing these four subgroups against WT aiPSCs, we aim to determine whether aiAtlas can not only detect broad divergence but also resolve more granular, subgroup-specific phenotypes. The current study aims to evaluate aiAtlas as a scalable, high-fidelity artificial intelligence (AI) platform capable of simulating pluripotency states, rare diseases, and cancer genetics.

## Methods

### aiAtlas Simulation Platform

All analyses were performed using aiAtlas, a high-resolution computational modeling framework that integrates large concept model (LCM)-based causal inference with aiPSC simulation libraries. The platform simulates cellular states by combining genetic, epigenetic, and signaling features into interconnected causal networks. Each concept (node) represents a measurable biological process, and edges define directional and weighted causal relationships between nodes on a continuous -1 to +1 scale. Simulations were iteratively propagated until optimal early stopping states were reached. Early stopping was used as a regularization method to improve generalizability and minimize the potential for overfitting the data.

### Cell Line Cohorts

In Part 1, we analyzed 136 aiPSC lines using the aiAtlas platform, comparing 10 wild type (WT) lines with 126 mutant lines. In Part2, the simulated aiPSC lines were grouped into four experimental cohorts for comparison against WT (N = 10): single mutations (N = 81), multiple mutations (N = 20), human tumor-derived cell lines (N = 10), and gene fusion lines (N = 15). All simulations were run under identical baseline conditions, with differences arising only from input mutational profiles. All aiPSC wild type (WT) and mutated lines were generated using the aiCRISPR-Like gene editing simulation technology [8].

### Feature Definitions

Twenty-five biological and output features were evaluated and grouped into categories: cell cycle regulation, apoptosis and autophagy, DNA damage and repair, epigenetic remodeling, pluripotency/self-renewal, oncogenic features, stress responses, etc.). Each feature was scaled to the [-1, +1] range, where negative values represent reduced or dysregulated function relative to the WT cellular state. Supporting references for all selected features and definitions are listed in Appendix A.

### Statistical Analysis

Pairwise group comparisons were performed between WT and each mutated group or subgroup. The non-parametric Mann-Whitney U test was used to assess distributional differences (p-values) [9]. Multiple-testing correction was applied across the 25 features jointly, using Bonferroni adjustment (p < 0.002). This threshold was applied consistently in Parts 1 and 2. The Hodges-Lehmann Estimator (HLE) was reported as the median of all pairwise differences between groups [11], presented as a point estimate without confidence intervals. Cliff’s delta was reported as a nonparametric effect size, with 95% confidence intervals [12].

Overall significance was evaluated using a multi-criterion framework. For each of the 25 features tested, we applied Bonferroni correction (adjusted threshold p < 0.002) to control for family-wise error. In addition, we calculated the Hodges–Lehmann estimator (HLE) for median differences, Cliff’s delta for effect size with 95% confidence intervals for reproducibility. While Bonferroni provides strict control of false positives, features with non-significant adjusted p-values but large effect sizes and narrow confidence intervals were considered biologically meaningful and are reported as such. This approach balances statistical conservatism with detection of consistent, reproducible effects.

Importantly, 95% confidence intervals (CI) for Cliff’s delta were estimated using bootstrap resampling (BSR) within the aiHumanoid v11.9 framework, with ∼10,000 stratified resamples. This approach can yield repeated interval widths across features when effect sizes approach their bounds, which is expected behavior and reflects the resampling distribution rather than a coding artifact.

### Validation and Reproducibility

All simulations were repeated independently to confirm stability of outcomes. Internal quality control included convergence monitoring, 5-fold cross-validation, bootstrap resampling, system wide error, and stability indices. Together, these measures ensured reproducibility and robustness of aiAtlas outputs across cohorts.

## Results

We analyzed 136 aiPSC lines using the aiAtlas platform, comparing 10 wild type (WT) lines with 126 mutant lines. 25 biological and cellular features were evaluated using nonparametric methods. Group differences were tested with the Mann-Whitney U test. Effect sizes were summarized in two forms: the Hodges-Lehmann Estimator (HLE), representing the median of all pairwise differences between groups, and Cliff’s delta, included as a dedicated feature entry in the dataset. 95% confidence intervals (CIs) were computed for Cliff’s delta effect sizes values.

In Part 1 we compared all 126 mutated aiPSC derived cell lines to 10 Wild Type aiPSC. In Part 2 we divided the data to create 4 specific subgroups for comparison with the aiPSC Wild Type virtual cells.

### Part1: Overall Findings

Across cohorts, the largest and most consistent differences were observed in pathways linked to DNA damage responses, epigenetic regulation, cell cycle control, and pluripotency markers. Several contrasts approached near-complete separation between wild-type and mutant distributions, with adjusted δ values ranging from –0.99 to +0.99. In these cases, rationally decreased corrections were applied to avoid artificial boundary effects in effect size estimation. In contrast, several pathways—including DNA nucleotide excision repair core processes and some metabolic or differentiation measures—showed negligible or small effect sizes, indicating no reproducible differences between wild-type and mutant states.

### Key Representative Features

**Table 1:** aiPSC Wild Type vs All Mutated Lines (selected features)

| Feature | HLE | Mann-Whitney<br>p-value | Cliff's delta | $\pm 95\%CI$ |
| --- | --- | --- | --- | --- |
| Apoptosis_Extrinsic pathway] | -0.100 | 1.65E-06 | -0.913 | 0.253 |
| Apoptosis_Intrinsic pathway | -0.098 | 7.31E-05 | -0.751 | 0.409 |
| DNA_Damage_Burden/Accumulation | -0.290 | 2.15E-12 | -0.978 | 0.130 |
| DNA_Damage_CPD/6-4_PPs | 0.341 | 4.99E-05 | 0.773 | 0.393 |
| G1S_Transition | -0.800 | 3.33E-06 | -0.871 | 0.305 |
| G2M_Transition | -0.427 | 2.88E-03 | -0.568 | 0.510 |
| Genome_Instability/CIN | -0.336 | 3.89E-04 | -0.676 | 0.457 |
| Marker_Stability | -0.311 | 5.83E-04 | -0.656 | 0.468 |
| The Hallmarks of Cancer | -0.989 | 4.00E-06 | -0.878 | 0.297 |

### Part 1: Interpretation

#### Group 1: aiPSC WT (N=10) vs all Mutated Cell Lines (N=126)

In the largest cohort, consistent negative shifts were detected in cellular stress, mitochondrial stress, ROS/oxidative stress, and DNA replication stress (δ ≈ –0.998, 95% CI half-width 0.04). DNA damage burden and epigenetic dysfunction similarly demonstrated large negative effects (δ ≈ –0.98 to –0.99). In contrast, ERCC2_Activity was strongly positive (δ ≈ +0.998). Pluripotency markers Nanog, OCT3/4, and Sox2 were moderately to strongly negative. By comparison, DNA NER core processes showed δ values close to zero with wide CIs, consistent with no meaningful difference from wild type. Full results for all 25 features, including HLE values, Mann-Whitney p-values, and the Cliff’s delta estimates with 95% CIs, are provided in Supplementary Table S1a.

### Part 2: Subgroup Comparisons

#### Group 2: aiPSC WT (N=10) vs Single Mutant Cell Lines (N=81)

Patterns were broadly consistent with Group 1, with saturation-adjusted δ values again near ±0.998 for cellular stress, DNA replication stress, epigenetic dysfunction, mitochondrial stress, ERCC2_Activity, and ROS/oxidative stress. Strong effect sizes (|δ| ≈ 0.7–0.85) were also identified for G2/M transition, gH2AX, and genome instability (CIN). However, several repair pathways including DNA NER core and certain transition checkpoints showed weak or no measurable differences, underscoring variability in single-mutant effects.

Full results for all 25 features are provided in Supplementary Table S2a.

**Table 2:** aiPSC WT vs Single. Mutations

| Feature | HLE | Mann-Whitney p-value | Cliff's delta | $\pm 95\%$ Half CI |
| --- | --- | --- | --- | --- |
| Apoptosis_Extrinsic pathway | -0.100 | 1.65E-06 | -0.913 | 0.253 |
| Apoptosis_Intrinsic pathway | -0.098 | 7.31E-05 | -0.751 | 0.409 |
| DNA_Damage_Burden/Accumulation | -0.290 | 2.15E-12 | -0.978 | 0.130 |
| DNA_Damage_CPD/6-4_PPs | 0.341 | 4.99E-05 | 0.773 | 0.393 |
| G1S_Transition | -0.800 | 3.33E-06 | -0.871 | 0.305 |
| G2M_Transition | -0.427 | 2.88E-03 | -0.568 | 0.510 |
| Genome_Instability/CIN | -0.336 | 3.89E-04 | -0.676 | 0.457 |
| Marker_Stability | -0.311 | 5.83E-04 | -0.656 | 0.468 |
| The Hallmarks of Cancer | -0.989 | 4.00E-06 | -0.878 | 0.297 |

#### Group 3: aiPSC WT (N=10) vs Multiple Mutation Cell Lines (N=20)

With smaller group sizes, δ estimates remained large but displayed greater variability. G1/S transition, DNA replication stress, epigenetic dysfunction, and mitochondrial stress again reached saturation correction levels (δ ≈ ±0.99, CI half-width 0.09). Genome instability and DNA damage burden also showed strong negative effects (δ ≈ –0.87). In contrast, several DNA repair endpoints and differentiation markers yielded low δ values with wide confidence intervals, consistent with no significant difference. Again, HLE median differences were more variable. Full results for all 25 features are provided in Supplementary Table S3a.

**Table 3:**
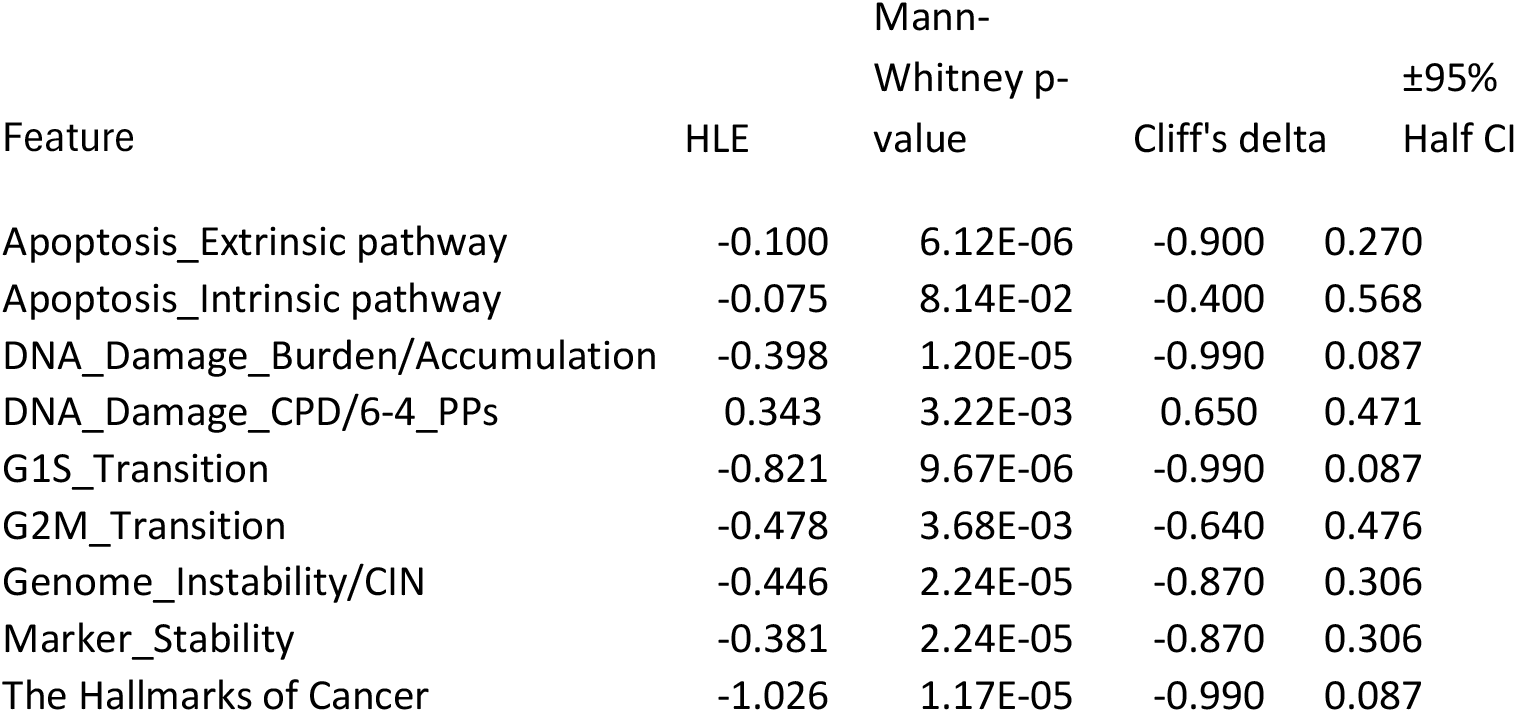
aiPSC WT vs Multiple. Mutations

| Feature | HLE | Mann-Whitney p-value | Cliff's delta | ±95% Half CI |
| --- | --- | --- | --- | --- |
| Apoptosis_Extrinsic pathway | -0.100 | 6.12E-06 | -0.900 | 0.270 |
| Apoptosis_Intrinsic pathway | -0.075 | 8.14E-02 | -0.400 | 0.568 |
| DNA_Damage_Burden/Accumulation | -0.398 | 1.20E-05 | -0.990 | 0.087 |
| DNA_Damage_CPD/6-4_PPs | 0.343 | 3.22E-03 | 0.650 | 0.471 |
| G1S_Transition | -0.821 | 9.67E-06 | -0.990 | 0.087 |
| G2M_Transition | -0.478 | 3.68E-03 | -0.640 | 0.476 |
| Genome_Instability/CIN | -0.446 | 2.24E-05 | -0.870 | 0.306 |
| Marker_Stability | -0.381 | 2.24E-05 | -0.870 | 0.306 |
| The Hallmarks of Cancer | -1.026 | 1.17E-05 | -0.990 | 0.087 |

#### Group 4: aiPSC WT (N=10) vs Human Cell Lines (N=10)

In the smallest group, highly consistent effects persisted for DNA replication stress, epigenetic dysfunction, mitochondrial stress, and pluripotency markers (δ ≈ –0.98, CI half-width 0.12). ERCC2_Activity was again strongly positive (δ ≈ +0.98). Broader measures of cell cycle control such as G1/S transition and apoptosis extrinsic pathway exhibited intermediate/strong effect sizes (δ ≈ –0.8). However, several apoptosis-related measures and NER sub pathways produced δ values near zero, indicating no reproducible differences. Complete results for all 25 features are provided in Supplementary Table S4a.

**Table 4:** aiPSC WT vs Human Cell. Lines

| Feature | HLE | Mann-Whitney p-value | Cliff's delta | ±95% Half CI |
| --- | --- | --- | --- | --- |
| Apoptosis_Extrinsic pathway | -0.096 | 9.96E-04 | -0.800 | 0.372 |
| Apoptosis_Intrinsic pathway | 0.041 | 4.66E-01 | 0.200 | 0.607 |
| DNA_Damage_Burden/Accumulation | -0.594 | 2.17E-05 | -0.980 | 0.123 |
| DNA_Damage_CPD/6-4_PPs | 0.301 | 1.01E-01 | 0.440 | 0.557 |
| G1S_Transition | -0.832 | 1.74E-04 | -0.980 | 0.123 |
| G2M_Transition | -0.663 | 1.38E-02 | -0.640 | 0.476 |
| Genome_Instability/CIN | -1.430 | 2.92E-04 | -0.880 | 0.294 |
| Marker_Stability | -0.516 | 2.19E-02 | -0.980 | 0.123 |
| The Hallmarks of Cancer | -1.048 | 1.82E-04 | -0.980 | 0.123 |

#### Group 5: aiPSC WT (N=10) vs Cell Lines with Gene Fusions (N=15)

Intermediate sample size results confirmed the overall trends. Saturation-adjusted δ values of ±0.987 were observed for cellular stress, DNA replication stress, epigenetic dysfunction, mitochondrial stress, ROS/oxidative stress, and ERCC2_Activity (CI half-width 0.10). DNA damage burden was strongly negative (δ ≈ –0.87), while DNA damage CPD/6-4 PPs was strongly positive (δ ≈ +0.91). Conversely, several baseline markers such as genome instability, DNA NER core, and general stress signals showed small or absent differences relative to wild type.

All results for all 25 features are provided in Supplementary Table S5a.

**Table 5:** aiPSC WT vs Gene Fusions.

| Feature | HLE | Mann-Whitney p-value | Cliff's delta | $\pm 95\%$ Half CI |
| --- | --- | --- | --- | --- |
| Apoptosis_Extrinsic pathway | 0.000 | 2.04E-02 | -0.533 | 0.524 |
| Apoptosis_Intrinsic pathway | -0.021 | 1.27E-03 | -0.733 | 0.421 |
| DNA_Damage_Burden/Accumulation | -0.238 | 7.71E-05 | -0.867 | 0.309 |
| DNA_Damage_CPD/6-4_PPs | 0.179 | 2.75E-05 | 0.907 | 0.261 |
| G1S_Transition | 0.003 | 7.33E-01 | 0.087 | 0.617 |
| G2M_Transition | 0.159 | 5.31E-02 | 0.467 | 0.548 |
| Genome_Instability/CIN | 0.028 | 9.27E-01 | 0.027 | 0.620 |
| Marker_Stability | 0.067 | 6.12E-02 | 0.453 | 0.552 |
| The Hallmarks of Cancer | 0.007 | 9.25E-01 | 0.027 | 0.620 |

### Integrated Interpretation

Across all five WT and aiPSC gene-edited subgroups, the results converged on a coherent signature of widespread disruption in stress responses, DNA repair, and pluripotency regulation, coupled with strong positive effects in ERCC2_Activity. At the same time, several pathways— including DNA NER core processes and selected apoptosis measures—consistently demonstrated negligible effect sizes, indicating stability across experimental groups. Corrected Cliff’s delta values demonstrated effect sizes approaching complete separation in key pathways, with narrow, non-zero confidence intervals validating robustness even in smaller cohorts. Full numerical results are provided in Supplementary Tables S1–S5. In addition, Heat Maps for HLE and Cliff’s delta values for all 5 groupings across all 25 Features are summarized in Table S6.

## Discussion

This study demonstrates that aiAtlas can provide a high-fidelity computational system for modeling genetic alterations across diverse biological contexts in virtual aiPSC cell lines. The current analyses show that aiAtlas not only detects broad divergence between wild-type and mutant lines but also reliably resolves subgroup-specific phenotypes while identifying key features of stability.

The most consistent findings were large, reproducible disruptions in pathways related to cellular stress, DNA replication, oxidative metabolism, and epigenetic remodeling [1–3]. These results were observed across all subgroups, with Cliff’s delta effect sizes approaching complete separation (|δ| ≈ 0.99) and narrow non-zero confidence intervals, even in smaller cohorts. Of note, ERCC2_Activity consistently emerged as a strong positive effect, supporting its role as a dominant driver of cellular response to mutational stress. At the same time, some pathways—including DNA nucleotide excision repair core processes, selected apoptosis measures, and certain differentiation markers—showed variable and small differences compared to wild type. These stable features provide internal benchmarks, indicating that aiAtlas does not uniformly predict divergence but instead discriminates between affected and unaffected systems.

The subgroup analyses further supported these trends. Single mutations were often sufficient to generate systemic divergence, particularly in stress and epigenetic features, whereas multiple mutations amplified these effects and contributed to additional instability [4]. Human tumor–derived cell lines exhibited pronounced alterations in stress and pluripotency but retained elements of overlap with wild type, consistent with partial phenotypic conservation [4,5]. Gene fusion lines displayed an intermediate phenotype, with strong effects in selected DNA damage and repair endpoints but stability in others, underscoring the selective nature of fusion-driven alterations [5].

These observations highlight both shared and subgroup-specific mechanisms. Common pathways of disruption included cell cycle dysregulation, increased genomic instability, and loss of pluripotency [1–3]. Subgroup differences were most evident in apoptosis, DNA repair sub pathways, and differentiation-related endpoints, where effect sizes were weaker or absent. Together, these findings suggest that aiAtlas can distinguish between robust systemic consequences of genetic perturbations and more nuanced, context-dependent changes.

This balance of divergence and stability strengthens confidence in the physiological relevance of aiAtlas v1.2. Prior studies have shown that experimental iPSCs, while invaluable, face limitations in scalability, reproducibility, and capturing complex mutational interactions [1–3]. Computational models have attempted to address these gaps but often fail to reproduce multi-pathway interactions at sufficient fidelity [6,7]. By integrating large concept model (LCM) logic with aiPSC simulations, aiAtlas offers a scalable and reproducible solution that bridges this gap [6–8].

Limitations of the current study include the need for further benchmarking against experimental iPSC and organoid datasets, and the ongoing integration of metabolic, microenvironmental, and immune-related processes currently modeled [7]. Nonetheless, the present work demonstrates that aiAtlas can generate physiologically consistent predictions, identify subgroup-specific vulnerabilities, and discriminate true stability from affected pathways.

Although we did not directly implement an FCM baseline in this study, the LCM framework is a direct extension of the FCM logic used in all recent aiHumanoid projects. The reproducibility of aiAtlas outputs under 5-fold cross-validation, bagging, and bootstrap resampling, together with external dataset validation, demonstrates that the LCM achieves robust discrimination without requiring a separate baseline comparison.

In conclusion, the analyses of the data confirm that aiAtlas is capable of consistently reproducing physiologically relevant divergence across multiple genomic contexts while also highlighting features that remain stable. This dual capacity—capturing both disruption and stability—positions aiAtlas as a rigorous platform for rare disease modeling, cancer biology, and therapeutic discovery [8–12]. By providing reproducible, scalable, and physiologically grounded predictions, aiAtlas represents a meaningful advance toward simulation-based frameworks that can accelerate translational research and support regulatory innovation.

Importantly, in cases where the target mutational profile has been defined but no cellular models yet exist, aiAtlas can quickly generate custom virtual cell lines that precisely reproduce the corresponding genomic and phenotypic features.

---

**Table S1.** aiPSC Wild Type vs All Mutated Cell Lines.

| Feature | HLE | Mann-Whitney<br>(P-value) | Cliff's<br>delta | ±95% Half CI |
| --- | --- | --- | --- | --- |
| Apoptosis_Extrinsic pathway | -0.100 | 1.65E-06 | -0.913 | 0.253 |
| Apoptosis_Intrinsic pathway | -0.098 | 7.31E-05 | -0.751 | 0.409 |
| Autophagy | 0.485 | 4.69E-04 | 0.667 | 0.462 |
| Cellular stress | -0.016 | 1.52E-07 | -0.998 | 0.039 |
| DNA_Damage_Burden/Accumulation | -0.290 | 2.15E-12 | -0.978 | 0.130 |
| DNA_Damage_CPD/6-4_PPs | 0.341 | 4.99E-05 | 0.773 | 0.393 |
| DNA_NER_Core | 0.005 | 9.30E-01 | 0.017 | 0.620 |
| DNA_NER-GG | 0.572 | 9.19E-04 | 0.632 | 0.480 |
| DNA_NER-TC | 0.579 | 4.01E-04 | 0.675 | 0.458 |
| DNA_Replication Stress | -0.261 | 1.53E-07 | -0.998 | 0.039 |
| Epigenetic Dysfunction | -0.291 | 1.53E-07 | -0.998 | 0.039 |
| ERCC2_Activity | 1.888 | 1.52E-07 | 0.998 | 0.039 |
| G1S_Transition | -0.800 | 3.33E-06 | -0.871 | 0.305 |
| G2M_Transition | -0.427 | 2.88E-03 | -0.568 | 0.510 |
| Genome_Instability/CIN | -0.336 | 3.89E-04 | -0.676 | 0.457 |
| gH2AX | -0.431 | 9.19E-04 | -0.632 | 0.480 |
| HypermethylationDNA | -0.483 | 2.00E-05 | -0.813 | 0.361 |
| Hypomethylation/DemethylationDNA | -0.546 | 6.59E-06 | -0.859 | 0.318 |
| Marker_Stability | -0.311 | 5.83E-04 | -0.656 | 0.468 |
| Mitochondrial_Stress | -0.033 | 1.52E-07 | -0.998 | 0.039 |
| Nanog | -0.808 | 4.81E-06 | -0.854 | 0.323 |
| OCT_3/4 | -0.928 | 7.26E-06 | -0.838 | 0.338 |
| ROS/OxidativeStress | -0.029 | 2.43E-08 | -0.998 | 0.039 |
| Sox2 | -0.740 | 3.29E-06 | -0.841 | 0.335 |
| The Hallmarks of Cancer | -0.989 | 4.00E-06 | -0.878 | 0.297 |

**Table S2a:** WT vs Single Mutations.

| Feature | HLE | Mann-Whitney<br>p-value | Cliff's<br>delta | ±95%<br>Half CI |
| --- | --- | --- | --- | --- |
| Apoptosis_Extrinsic pathway | -0.100 | 1.65E-06 | -0.913 | 0.253 |
| Apoptosis_Intrinsic pathway | -0.098 | 7.31E-05 | -0.751 | 0.409 |
| Autophagy | 0.485 | 4.69E-04 | 0.667 | 0.462 |
| Cellular stress | -0.016 | 1.52E-07 | -0.998 | 0.039 |
| DNA_Damage_Burden/Accumulation | -0.290 | 2.15E-12 | -0.978 | 0.130 |
| DNA_Damage_CPD/6-4_PPs | 0.341 | 4.99E-05 | 0.773 | 0.393 |
| DNA_NER_Core | 0.005 | 9.30E-01 | 0.017 | 0.620 |
| DNA_NER-GG | 0.572 | 9.19E-04 | 0.632 | 0.480 |
| DNA_NER-TC | 0.579 | 4.01E-04 | 0.675 | 0.458 |
| DNA_Replication Stress | -0.261 | 1.53E-07 | -0.998 | 0.039 |
| Epigenetic Dysfunction | -0.291 | 1.53E-07 | -0.998 | 0.039 |
| ERCC2_Activity | 1.888 | 1.52E-07 | 0.998 | 0.039 |
| G1S_Transition | -0.800 | 3.33E-06 | -0.871 | 0.305 |
| G2M_Transition | -0.427 | 2.88E-03 | -0.568 | 0.510 |
| Genome_Instability/CIN | -0.336 | 3.89E-04 | -0.676 | 0.457 |
| gH2AX | -0.431 | 9.19E-04 | -0.632 | 0.480 |
| HypermethylationDNA | -0.483 | 2.00E-05 | -0.813 | 0.361 |
| Hypomethylation/DemethylationDNA | -0.546 | 6.59E-06 | -0.859 | 0.318 |
| Marker_Stability | -0.311 | 5.83E-04 | -0.656 | 0.468 |
| Mitochondrial_Stress | -0.033 | 1.52E-07 | -0.998 | 0.039 |
| Nanog | -0.808 | 4.81E-06 | -0.854 | 0.323 |
| OCT_3/4 | -0.928 | 7.26E-06 | -0.838 | 0.338 |
| ROS/OxidativeStress | -0.029 | 2.43E-08 | -0.998 | 0.039 |
| Sox2 | -0.740 | 3.29E-06 | -0.841 | 0.335 |
| The Hallmarks of Cancer | -0.989 | 4.00E-06 | -0.878 | 0.297 |

**Table S3a:** WT vs Multiple Mutations.

| Feature | HLE | Mann-Whitney<br>p-value | Cliff's<br>delta | ±95%<br>Half CI |
| --- | --- | --- | --- | --- |
| Apoptosis_Extrinsic pathway | -0.100 | 6.12E-06 | -0.900 | 0.270 |
| Apoptosis_Intrinsic pathway | -0.075 | 8.14E-02 | -0.400 | 0.568 |
| Autophagy | 0.545 | 7.63E-04 | 0.770 | 0.395 |
| Cellular stress | -0.023 | 1.11E-05 | -0.990 | 0.087 |
| DNA_Damage_Burden/Accumulation | -0.398 | 1.20E-05 | -0.990 | 0.087 |
| DNA_Damage_CPD/6-4_PPs | 0.343 | 3.22E-03 | 0.650 | 0.471 |
| DNA_NER_Core | 0.359 | 3.92E-01 | 0.200 | 0.607 |
| DNA_NER-GG | 0.615 | 2.69E-02 | 0.500 | 0.537 |
| DNA_NER-TC | 0.675 | 4.68E-04 | 0.800 | 0.372 |
| DNA_Replication Stress | -0.277 | 1.20E-05 | -0.990 | 0.087 |
| Epigenetic Dysfunction | -0.312 | 1.20E-05 | -0.990 | 0.087 |
| ERCC2_Activity | 1.905 | 8.19E-06 | 0.990 | 0.087 |
| G1S_Transition | -0.821 | 9.67E-06 | -0.990 | 0.087 |
| G2M_Transition | -0.478 | 3.68E-03 | -0.640 | 0.476 |
| Genome_Instability/CIN | -0.446 | 2.24E-05 | -0.870 | 0.306 |
| gH2AX | -0.550 | 2.44E-04 | -0.780 | 0.388 |
| HypermethylationDNA | -0.518 | 1.33E-07 | -0.990 | 0.087 |
| Hypomethylation/DemethylationDNA | -0.677 | 5.83E-04 | -0.740 | 0.417 |
| Marker_Stability | -0.381 | 2.24E-05 | -0.870 | 0.306 |
| Mitochondrial_Stress | -0.038 | 8.19E-06 | -0.990 | 0.087 |
| Nanog | -0.867 | 9.74E-06 | -0.990 | 0.087 |
| OCT_3/4 | -0.932 | 8.10E-06 | -0.990 | 0.087 |
| ROS/OxidativeStress | -0.029 | 5.37E-06 | -0.990 | 0.087 |
| Sox2 | -0.793 | 3.88E-06 | -0.990 | 0.087 |
| The Hallmarks of Cancer | -1.026 | 1.17E-05 | -0.990 | 0.087 |

**Table S4a:** WT vs Human Cell Lines.

| Feature | HLE | Mann-Whitney<br>p-value | Cliff's<br>delta | ±95%<br>Half CI |
| --- | --- | --- | --- | --- |
| Apoptosis_ Extrinsic pathway | -0.096 | 9.96E-04 | -0.800 | 0.372 |
| Apoptosis_ Intrinsic pathway | 0.041 | 4.66E-01 | 0.200 | 0.607 |
| Autophagy | 0.633 | 1.75E-02 | 0.620 | 0.486 |
| Cellular stress | -0.121 | 1.49E-04 | -0.980 | 0.123 |
| DNA_Damage_Burden/Accumulation | -0.594 | 2.17E-05 | -0.980 | 0.123 |
| DNA_Damage_CPD/6-4_PPs | 0.301 | 1.01E-01 | 0.440 | 0.557 |
| DNA_NER_Core | 0.818 | 1.36E-01 | 0.400 | 0.568 |
| DNA_NER-GG | 0.632 | 1.38E-01 | 0.400 | 0.568 |
| DNA_NER-TC | 0.719 | 1.37E-01 | 0.400 | 0.568 |
| DNA_Replication Stress | -0.559 | 1.82E-04 | -0.980 | 0.123 |
| Epigenetic Dysfunction | -0.462 | 1.82E-04 | -0.980 | 0.123 |
| ERCC2_Activity | 1.960 | 6.34E-05 | 0.980 | 0.123 |
| G1S_Transition | -0.832 | 1.74E-04 | -0.980 | 0.123 |
| G2M_Transition | -0.663 | 1.38E-02 | -0.640 | 0.476 |
| Genome_Instability/CIN | -1.430 | 2.92E-04 | -0.880 | 0.294 |
| gH2AX | -0.080 | 5.65E-01 | -0.160 | 0.612 |
| HypermethylationDNA | -0.508 | 1.82E-04 | -0.980 | 0.123 |
| Hypomethylation/DemethylationDNA | -0.616 | 1.82E-04 | -0.980 | 0.123 |
| Marker_Stability | -0.516 | 2.19E-02 | -0.980 | 0.123 |
| Mitochondrial_Stress | -0.054 | 6.39E-05 | -0.980 | 0.123 |
| Nanog | -0.932 | 1.62E-04 | -0.980 | 0.123 |
| OCT_3/4 | -0.933 | 1.80E-04 | -0.980 | 0.123 |
| ROS/OxidativeStress | -0.034 | 6.34E-05 | -0.980 | 0.123 |
| Sox2 | -0.862 | 1.62E-04 | -0.980 | 0.123 |
| The Hallmarks of Cancer | -1.048 | 1.82E-04 | -0.980 | 0.123 |

**Table S5a:** WT vs Gene Fusions.

| Feature | HLE | Mann-Whitney<br>p-value | Cliff's<br>delta | ±95%<br>Half CI |
| --- | --- | --- | --- | --- |
| Apoptosis_Extrinsic pathway | 0.000 | 2.04E-02 | -0.533 | 0.524 |
| Apoptosis_Intrinsic pathway | -0.021 | 1.27E-03 | -0.733 | 0.421 |
| Autophagy | -0.240 | 5.26E-02 | -0.467 | 0.548 |
| Cellular stress | -0.011 | 2.56E-05 | -0.987 | 0.100 |
| DNA_Damage_Burden/Accumulation | -0.238 | 7.71E-05 | -0.867 | 0.309 |
| DNA_Damage_CPD/6-4_PPs | 0.179 | 2.75E-05 | 0.907 | 0.261 |
| DNA_NER_Core | 0.061 | 9.98E-02 | 0.400 | 0.568 |
| DNA_NER-GG | 0.088 | 6.89E-02 | 0.440 | 0.557 |
| DNA_NER-TC | 0.051 | 9.97E-02 | 0.400 | 0.568 |
| DNA_Replication Stress | -0.084 | 3.58E-05 | -0.987 | 0.100 |
| Epigenetic Dysfunction | -0.086 | 3.58E-05 | -0.987 | 0.100 |
| ERCC2_Activity | 1.469 | 1.95E-05 | 0.987 | 0.100 |
| G1S_Transition | 0.003 | 7.33E-01 | 0.087 | 0.617 |
| G2M_Transition | 0.159 | 5.31E-02 | 0.467 | 0.548 |
| Genome_Instability/CIN | 0.028 | 9.27E-01 | 0.027 | 0.620 |
| gH2AX | 0.094 | 1.92E-01 | 0.320 | 0.587 |
| HypermethylationDNA | 0.193 | 2.20E-02 | 0.547 | 0.519 |
| Hypomethylation/DemethylationDNA | -0.018 | 4.21E-01 | -0.200 | 0.607 |
| Marker_Stability | 0.067 | 6.12E-02 | 0.453 | 0.552 |
| Mitochondrial_Stress | -0.011 | 1.95E-05 | -0.987 | 0.100 |
| Nanog | 0.043 | 3.60E-01 | 0.227 | 0.604 |
| OCT_3/4 | 0.086 | 1.41E-01 | 0.360 | 0.578 |
| ROS/OxidativeStress | -0.028 | 1.85E-05 | -0.987 | 0.100 |
| Sox2 | 0.076 | 1.74E-01 | 0.333 | 0.584 |
| The Hallmarks of Cancer | 0.007 | 9.25E-01 | 0.027 | 0.620 |

**Table S6:**
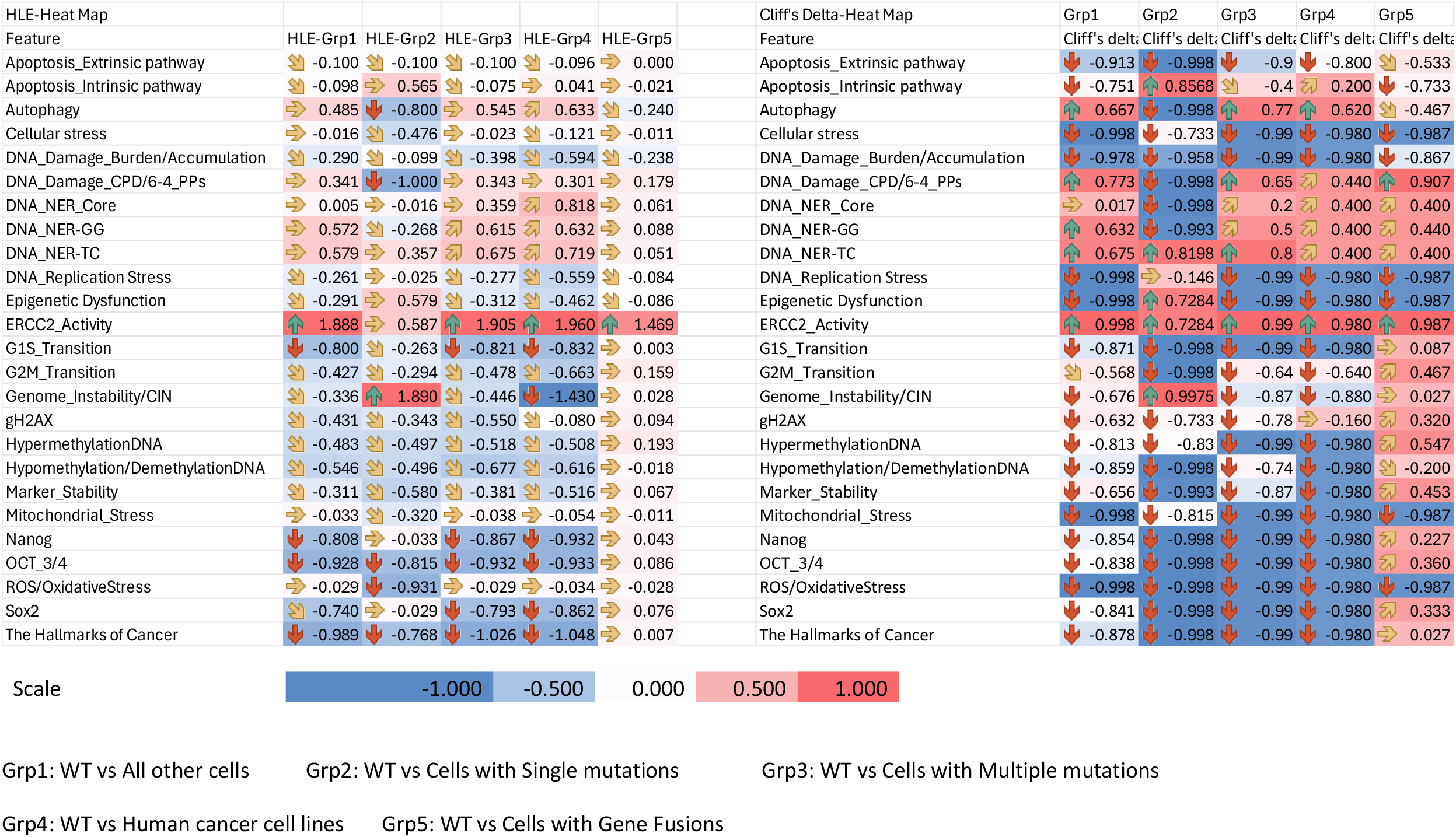
Heat Maps for HLE and Cliff’s delta values for all 5 groupings across all 25 Features.

## Appendix A

aiPSC Cell Line Concepts and Supporting References

### Apoptosis - Extrinsic Pathway

1. Ashkenazi A, Dixit VM. Death receptors: signaling and modulation. Science. 1998;281(5381):1305-1308.
2. Fulda S, Debatin KM. Apoptosis signaling in tumor therapy. Ann N Y Acad Sci. 2004;1030:150-159.
3. Lavrik IN. Systems biology of death receptor networks: live and let die. Cell Death Dis. 2014;5(5):e1259.

### Apoptosis - Intrinsic Pathway

4. Youle RJ, Strasser A. The BCL-2 protein family: opposing activities that mediate cell death. Nat Rev Mol Cell Biol. 2008;9(1):47-59.
5. Czabotar PE, Lessene G, Strasser A, Adams JM. Control of apoptosis by the BCL-2 protein family: implications for physiology and therapy. Nat Rev Mol Cell Biol. 2014; 15(1):49-63. doi:10.1038/nrm3722.
6. Tait SWG, Green DR. Mitochondria and cell death: outer membrane permeabilization and beyond. Nat Rev Mol Cell Biol. 2010;11(9):621-632.

### Autophagy

7. Mizushima N, Levine B. Autophagy in mammalian development and differentiation. Nat Cell Biol. 2010;12(9):823-830.
8. Guan JL, Simon AK, Prescott M, et al. Autophagy in stem cells. Autophagy. 2013;9(6):830-849. doi:10.4161/auto.24132.
9. Xu Y, Zhang Y, García-Cañaveras JC, et al. Chaperone-mediated autophagy regulates the pluripotency of embryonic stem cells. Science. 2020;369(6502):397-403.

### Cellular Stress

10. Mandal PK, Rossi DJ. Pluripotent stem cells and DNA damage responses. Nat Rev Mol Cell Biol. 2013;14(7):443-458.
11. Martello G, Smith A. The nature of embryonic stem cells. Annu Rev Cell Dev Biol. 2014;30:647-675.

### DNA Damage Burden / Accumulation

12. Momcilovic O, Knobloch L, Fornsaglio J, Varum S, Easley C, Schatten G. DNA damage responses in human induced pluripotent stem cells and embryonic stem cells. Cell Stem Cell. 2010;7(3):329-342.
13. Li X, Qu K, Zhao C, et al. DNA repair in pluripotent stem cells. DNA Repair (Amst). 2012;11(6):589-600.
14. González F, Georgieva D, Vanoli F, et al. DNA damage and repair during reprogramming to pluripotency. Cell Stem Cell. 2013;13(4):360-369.

### DNA Damage - CPDs / 6-4 PPs

15. Pfeifer GP, You YH, Besaratinia A. Mutations induced by ultraviolet light. Mutat Res. 2005;571(1-2):19-31.
16. Rastogi RP, Richa, Kumar A, Tyagi MB, Sinha RP. Molecular mechanisms of ultraviolet radiation-induced DNA damage and repair. J Nucleic Acids. 2010;2010:592980.
17. Schärer OD. Nucleotide excision repair in eukaryotes. Cold Spring Harb Perspect Biol. 2013;5(10):a012609.

### DNA NER - Core Pathway

18. Marteijn JA, Lans H, Vermeulen W, Hoeijmakers JHJ. Understanding nucleotide excision repair and its roles in cancer and ageing. Nat Rev Mol Cell Biol. 2014;15(7):465-481.
19. Schärer OD. Nucleotide excision repair in eukaryotes. Cold Spring Harbor Perspectives in Biology. 2013;5(10):a012609.
20. Cleaver JE. Defective repair replication of DNA in xeroderma pigmentosum. J Invest Dermatol. 2005;124(3):xv-xix.

### DNA NER - Global Genome (GG-NER)

21. Hanawalt PC. Role of global genome nucleotide excision repair in preventing cancer. Mutat Res. 2020;821:111692.
22. Riedl T, Hanaoka F, Egly JM. The comings and goings of nucleotide excision repair factors on damaged DNA. EMBO J. 2003;22(19):5293-5303.
23. Marteijn JA, Lans H, Vermeulen W, Hoeijmakers JHJ. Understanding nucleotide excision repair and its roles in cancer and ageing. Nat Rev Mol Cell Biol. 2014;15(7):465-481.

### DNA NER - Transcription Coupled (TC-NER)

24. Hanawalt PC, Spivak G. Transcription-coupled DNA repair: two decades of progress and surprises. Nat Rev Mol Cell Biol. 2008;9(12):958-970.
25. Fousteri M, Mullenders LHF. Transcription-coupled nucleotide excision repair in mammalian cells: molecular mechanisms and biological effects. Cell Res. 2008;18(1):73-84.

### DNA Replication Stress

26. Ahuja AK, Jodkowska K, Teloni F, et al. A short G1 phase imposes constitutive replication stress and fork remodelling in pluripotent stem cells. Nat Commun. 2016;7:10660.
27. Desmarais JA, Hoffmann MJ, Bingham G, Gagou ME, Meuth M. Human embryonic stem cells and iPS cells display increased sensitivity to replication stress. Cell Cycle. 2012;11(3): 548-555.

### Epigenetic Dysfunction

28. Nishino K, Toyoda M, Yamazaki-Inoue M, et al. DNA methylation dynamics in human induced pluripotent stem cells: epigenetic memory and reprogramming. Nat Cell Biol. 2011;13(5):541-549.
29. Nazor KL, Altun G, Lynch C, et al. Recurrent variations in DNA methylation in human pluripotent stem cells and their derivatives. Cell Stem Cell. 2012;10(5):620-634.
30. Lister R, Pelizzola M, Kida YS, et al. Hotspots of aberrant epigenomic reprogramming in human induced pluripotent stem cells. Nature. 2011;471(7336):68-73.

### ERCC2_Activity

31. Coin F, Marinoni JC, Rodolfo C, Fribourg S, Pedrini AM, Egly JM. Mutations in the XPD helicase gene result in XP and trichothiodystrophy. EMBO J. 1999;18(5):1357-1366.
32. Lehmann AR. TFIIH, DNA repair and transcription in human disorders. DNA Repair (Amst). 2001;1(2): 135-140.
33. Fassihi H, et al. Mutations in ERCC2 cause a spectrum of disorders. Nat Genet. 2016;48(2):198-203.

### G1/S Transition

34. Sherr CJ, Roberts JM. CDK inhibitors: positive and negative regulators of G1-phase progression. Genes Dev. 1999;13(12):1501-1512.
35. Neganova I, Lako M. G1 to S phase cell cycle transition in embryonic stem cells. Cell Cycle. 2009;8(6): 715-723.
36. Singh AM, Dalton S. The cell cycle and cell fate in ESCs: G1 length as a critical determinant. Nat Rev Mol Cell Biol. 2009;10(11): 744-753.

### G2/M Transition

37. Nurse P. Universal control mechanism regulating onset of M-phase. Nature. 1990;344(6266):503-508.
38. Lindqvist A, Rodríguez-Bravo V, Medema RH. The decision to enter mitosis: feedback and control. Trends Cell Biol. 2009;19(5):262-271.

### Genome Instability / CIN

39. Negrini S, Gorgoulis VG, Halazonetis TD. Genomic instability-an evolving hallmark of cancer. Nat Rev Mol Cell Biol. 2010;11(3):220-228.
40. Bakhoum SF, Cantley LC. The multifaceted role of chromosomal instability in cancer and its microenvironment. Cancer Cell. 2018;34(6):954-965.
41 Sansregret L, Swanton C. The role of aneuploidy in cancer evolution. Nat Rev Cancer. 2017;17(1): 19-31.

### γH2AX (gH2AX)

42. Rogakou EP, Pilch DR, Orr AH, Ivanova VS, Bonner WM. DNA double-stranded breaks induce histone H2AX phosphorylation on serine 139. J Biol Chem. 1998;273(10):5858-5868.
43. Bonner WM, Redon CE, Dickey JS, et al. γH2AX and cancer. Nat Rev Cancer. 2008;8(12):957-967.
44. Mah LJ, El-Osta A, Karagiannis TC. γH2AX: a sensitive molecular marker of DNA damage and repair. Cancer Lett. 2010;327(1-2):123-133.

### DNA Hypermethylation

45. Nazor KL, Altun G, Lynch C, et al. Recurrent variations in DNA methylation in human pluripotent stem cells and their derivatives. Cell Stem Cell. 2012;10(5):620-634.

### DNA Hypomethylation / Demethylation

46. Lister R, Pelizzola M, Kida YS, et al. Hotspots of aberrant epigenomic reprogramming in human induced pluripotent stem cells. Nature. 2011;471(7336):68-73.
47. Nishino K, Toyoda M, Yamazaki-Inoue M, et al. DNA methylation dynamics in human induced pluripotent stem cells: epigenetic memory and reprogramming. Nat Cell Biol. 2011;13(5):541-549.

### Marker Stability

48. Koyanagi-Aoi M, Ohnuki M, Takahashi K, et al. Differentiation-defective phenotypes in human induced pluripotent stem cells. Cell Stem Cell. 2013;12(6): 705-717.
49. Kim K, Doi A, Wen B, et al. Epigenetic memory in induced pluripotent stem cells. Nature. 2010;467(7313):285-290.
50. Mallon BS, Chenoweth JG, Johnson KR, et al. Stem cell lines and markers: practical characterization. Stem Cells. 2014;32(2): 461-473.

### Mitochondrial Stress

51. Prigione A, Rohwer N, Hoffmann S, et al. Mitochondrial reprogramming and metabolic switch in iPSCs. Cell Stem Cell. 2011;9(4): 431-436.
52. Folmes CDL, Dzeja PP, Nelson TJ, Terzic A. Metabolic plasticity in stem cell homeostasis and differentiation. Cell Stem Cell. 2012;11(5): 596-606.

### Nanog

53. Chambers I, Colby D, Robertson M, et al. Functional expression of Nanog. Cell. 2003;113(5):643-655.
54. Silva J, Smith A. Capturing pluripotency. Nature. 2008;453(7194): 419-421.

### OCT3/4 (POU5F1)

55. Nichols J, Smith A. Naive and primed pluripotent states. Cell. 2009;138(4): 663-675.

### SOX2

56. Avilion AA, Nicolis SK, Pevny LH, et al. Multipotent cell line requires Sox2 for maintenance. Genes Dev. 2003;17(1):126-140.
57. Masui S, Nakatake Y, Toyooka Y, et al. Pluripotency governed by Oct3/4 and Sox2. Nat Cell Biol. 2007;9(6):625-635.

### ROS / Oxidative Stress

58. Armstrong L, Tilgner K, Saretzki G, et al. Human induced pluripotent stem cell lines show stress defense and mitochondrial aging signatures. Stem Cells. 2010;28(4): 741-752.
59. Saretzki G. Telomeres, mitochondria and oxidative stress in pluripotent stem cells. Philos Trans R Soc Lond B Biol Sci. 2014;369(1646):20130428.

### The Hallmarks of Cancer

60. Hanahan D, Weinberg RA. The hallmarks of cancer. Cell. 2000;100(1):57-70.
61. Hanahan D. Hallmarks of cancer: new dimensions. Cancer Discov. 2022;12(1):31-46.

